# Covid-19 vaccine immunogenicity in people living with HIV-1

**DOI:** 10.1101/2021.08.13.456258

**Authors:** Lauriane Nault, Lorie Marchitto, Guillaume Goyette, Daniel Tremblay-Sher, Claude Fortin, Valérie Martel-Laferrière, Benoît Trottier, Jonathan Richard, Madeleine Durand, Daniel Kaufmann, Andrés Finzi, Cécile Tremblay

## Abstract

**Introduction:** COVID-19 vaccine efficacy has been evaluated in large clinical trials and in real-world situation. Although they have proven to be very effective in the general population, little is known about their efficacy in immunocompromised patients. HIV-infected individuals’ response to vaccine may vary according to the type of vaccine and their level of immunosuppression. We evaluated immunogenicity of an mRNA anti-SARS CoV-2 vaccine in HIV-positive individuals.

**Methods:** HIV-positive individuals (n=121) were recruited from HIV clinics in Montreal and stratified according to their CD4 counts. A control group of 20 health care workers naïve to SARS CoV-2 was used. The participants’ Anti-RBD IgG responses were measured by ELISA at baseline and 3 to 4 weeks after receiving the first dose of an mRNA vaccine).

**Results:** Eleven of 121 participants had anti-COVID-19 antibodies at baseline, and a further 4 had incomplete data for the analysis. Mean anti-RBD IgG responses were similar between between the HIV negative control group (n=20) and the combined HIV+ group (n=106) (p = 0.72). However, these responses were significantly lower in the group with <250 CD4 cells/mm^3^. (p<0.0001). Increasing age was independently associated with decreased immunogenicity.

**Conclusion:** HIV-positive individuals with CD4 counts over 250 cells/mm^3^ have an anti-RBD IgG response similar to the general population. However, HIV-positive individuals with the lowest CD4 counts (<250 cells/mm^3^) have a weaker response. These data would support the hypothesis that a booster dose might be needed in this subgroup of HIV-positive individuals, depending on their response to the second dose.

---

Anti-COVID-19 vaccines have been developed at an extraordinary pace and have proven to be extremely effective in clinical trials^1–3^ and in a real-world setting^4,5^. However, these studies provided little information on vaccine immunogenicity in immunocompromised individuals. Several studies have evaluated anti-COVID-19 vaccine responses in immunocompromised patients, mostly in transplant patients or patients with auto-immune disease, cancer, or on dialysis^6–14^. They showed a range of antibody responses from 14% in solid organ transplant recipient to 57-96% in hemodialysis patients^14^. Few studies have addressed vaccine immunogenicity in HIV-infected individuals^15–18^. We seek to evaluate immunogenicity of an mRNA anti-SARS CoV-2 vaccine in HIV-positive individuals, and determine the impact of CD4 T cell counts on vaccine response.

## Methods

### Study population and design

121 HIV-positive individuals treated with antiretroviral therapy were recruited from HIV clinics in Montreal and stratified according to their CD4 counts (<250 cells/mm3, between 250 and 500 cells/mm3, and >500 cells/mm3). A control group of 20 health care workers naïve to COVID-19 was used. The participants’ immunogenicity was measured at baseline and between 3 and 4 weeks after receiving the first dose of an mRNA vaccine (Moderna mRNA-1273 in the HIV+ subgroups, Pfizer BNT162b2 in the HIV-control group).

### Antibody measurement

#### Plasmid

The plasmid expressing SARS-CoV-2 S RBD was previously reported^19^.

#### Protein expression and purification

FreeStyle 293F cells (Invitrogen) were grown in FreeStyle 293F medium (Invitrogen) to a density of 1 × 10^6^ cells/mL at 37°C with 8 % CO2 with regular agitation (150 rpm). Cells were transfected with a plasmid coding for SARS-CoV-2 S RBD (using ExpiFectamine 293 transfection reagent), as directed by the manufacturer (Invitrogen). One week later, cells were pelleted and discarded. Supernatants were filtered using a 0.22 μm filter (Thermo Fisher Scientific). Recombinant RBD was purified by nickel affinity column, as directed by the manufacturer (Invitrogen). The RBD preparations were dialyzed against phosphate-buffered saline (PBS) and stored in aliquots at −80°C until further use. To assess purity, recombinant proteins were loaded on SDS-PAGE gels and stained with Coomassie Blue.

#### Plasma and antibodies

Plasma samples were heat-inactivated for 1 hour at 56 °C and stored at −80°C until ready to use in subsequent experiments. Plasma from uninfected donors collected before the pandemic were used as negative controls and used to calculate the seropositivity threshold in our ELISA assay. The monoclonal antibody CR3022 was used as a positive control in the ELISA assay and was previously described^14,19–23^. Horseradish peroxidase (HRP)-conjugated antibody specific for the Fc region of human IgG (Invitrogen) was used as secondary antibody to detect antibody binding in ELISA experiments.

#### ELISA

Anti-RBD IgG responses were measured by ELISA 4 weeks after the first dose. The SARS-CoV-2 RBD assay used was previously described ^19,22^. Briefly, recombinant SARS-CoV-2 S RBD (2.5 μg/ml), or bovine serum albumin (BSA) (2.5 μg/ml) as a negative control, were prepared in PBS and were adsorbed to plates (MaxiSorp Nunc) overnight at 4°C. Coated wells were subsequently blocked with blocking buffer (Tris-buffered saline [TBS] containing 0.1% Tween20 and 2% BSA) for 1h at room temperature.

Wells were then washed four times with washing buffer (Tris-buffered saline [TBS] containing 0.1% Tween20). CR3022 mAb (50ng/ml) or human plasma (1/500) were prepared in a diluted solution of blocking buffer (0.1 % BSA) and incubated with the RBD-coated wells for 90 minutes at room temperature. Plates were washed four times with washing buffer followed by incubation with secondary Abs (diluted in a diluted solution of blocking buffer (0.4% BSA)) for 1h at room temperature, followed by four washes. HRP enzyme activity was determined after the addition of a 1:1 mix of Western Lightning oxidizing and luminol reagents (Perkin Elmer Life Sciences). Light emission was measured with a LB941 TriStar luminometer (Berthold Technologies). Signal obtained with BSA was subtracted for each plasma and was then normalized to the signal obtained with CR3022 mAb present in each plate. The seropositivity threshold was established using the following formula: mean of all pre-pandemic plasma + (3 standard deviation of the mean of all pre-pandemic plasma).

### Statistical analysis

Anti-RBD IgG values were log transformed for the analysis. The HIV-negative control group^23^ and HIV-infected combined group were compared with a two-tailed t test for anti-RBD titer, and a chi-square test for proportions of individuals reaching measurable anti-RBD antibodies. The association between age, sex and levels of CD4 was assessed using uni and multivariable linear regression models, with factors showing an association in the univariable models included into the multivariable model. Tukey tests were used for between group comparisons. Immune compromise was described categorically (CD4 count <250, between 250 and 500, above 500, and HIV-control). Age was integrated as a continuous variable. Type 3 sums of squares were used to account for design imbalance (small number of participants in the CD4<250 group). No significant interaction was detected between the independent variables. Statistical analysis was conducted using R version 4.1.

## Results

We present the immunogenicity results at week 3-4 after the participants’ first vaccine dose. Participants’ characteristics are described in Table 1. Eleven of 121 participants had anti-COVID-19 antibodies at baseline, suggesting prior exposure to COVID-19, and were excluded from the analysis. Four additional participants had incomplete CD4+ count information and were not included in the analysis. There were no statistically significant differences in immunogenicity between the HIV-control group (n=20) and the combined HIV+ group (n=106) either in magnitude (difference of means, two tailed t test, p = 0.72) or in the proportion of individuals mounting a measurable immune response (HIV-: 19/20 (95%) vs HIV+: 100/106 (94.3%), p = 0.91). Results from the multivariable linear regression, showing the associations between CD4 levels, age and anti-RBD antibody titers, are presented in Table 2. Both CD4 stratification and age were significantly associated with immunogenicity. Between group comparisons show that mean anti-RBD IgG responses were significantly lower in the CD4<250 group compared to all other groups, independent of age (p<0.001) (Figure 1). The mean relative luminescence units (RLU) for anti-RBD antibodies was 4.75 in participants with CD4<250, compared to 50.71 for the remainder of the study sample. There were no significant differences in immunogenicity among other groups (CD4>250 or HIV negative.) Independently, age was also significantly, but weakly, associated with decreased immunogenicity. For every increase of 10 years in age, the model predicted a decrease of 1.33 RLU. The range of ELISA RLU in this population was 2.56 (detection limit) to 236.03). Sex was not associated with immunogenicity. Although the regression model fit was significant (p < 0.00001), the adjusted R squared was only 0.24, meaning that CD4 counts and age combined only account for a relatively small proportion of the variance in immunogenicity in the study population (Figure 2).

**Table 1:** Participant characteristics

| Participant characteristics | Age<br>mean [range] | Sex<br>Male (%) |
| --- | --- | --- |
| HIV- Controls (n=20) | 47 [21, 59] | 7 (35.0%) |
| HIV+ Combined (n=106) | 43 [21, 65] | 90 (84.9%) |
| CD4<250 (n=6) | 48 [24, 61] | 5 (86.3%) |
| 250<CD4<500 (n=18) | 49 [34, 60] | 15 (88.9%) |
| CD4>500 (n=82) | 41 [21, 65] | 69 (84.1%) |

**Table 2:** Uni- and multi-variable regression models. Immunogenicity (the dependent variable) was log transformed for the analysis. Sex was not found to be significantly associated in the univariate model and was not included in the multivariate model. No significant interaction was detected between the age and stratification variables (not shown).

| Variable | Univariable models<br>Beta coefficient<br>[95% CI] | p-value | Multivariable models<br>Beta coefficient<br>[95% CI] | p-value |
| --- | --- | --- | --- | --- |
| Age (per 10 year increase in age) | -0.317<br>[-0.489, -0.144] | <0.001 | -0.289<br>[-0.456, -0.121] | <0.001 |
| Sex (male vs female) | 0.289<br>[-0.732, 0.154] | 0.199 |  |  |
| CD4 stratification (each group compared to the entire study population) |  |  |  |  |
| <250 | -1.593<br>[-2.199, -0.987] | <0.0001 | -1.547<br>[-2.129, -0.965] | <0.0001 |
| 250-500 | 0.439<br>[0.032, 0.846] | 0.035 | 0.536<br>[0.141, 0.930] | 0.008 |
| >500 | 0.608<br>[0.313, 0.902] | <0.001 | 0.453<br>[0.156, 0.749] | 0.003 |
| HIV- | 0.546<br>[0.152, 0.941] | 0.007 | 0.558<br>[0.180, 0.937] | 0.004 |

**Figure 1:**
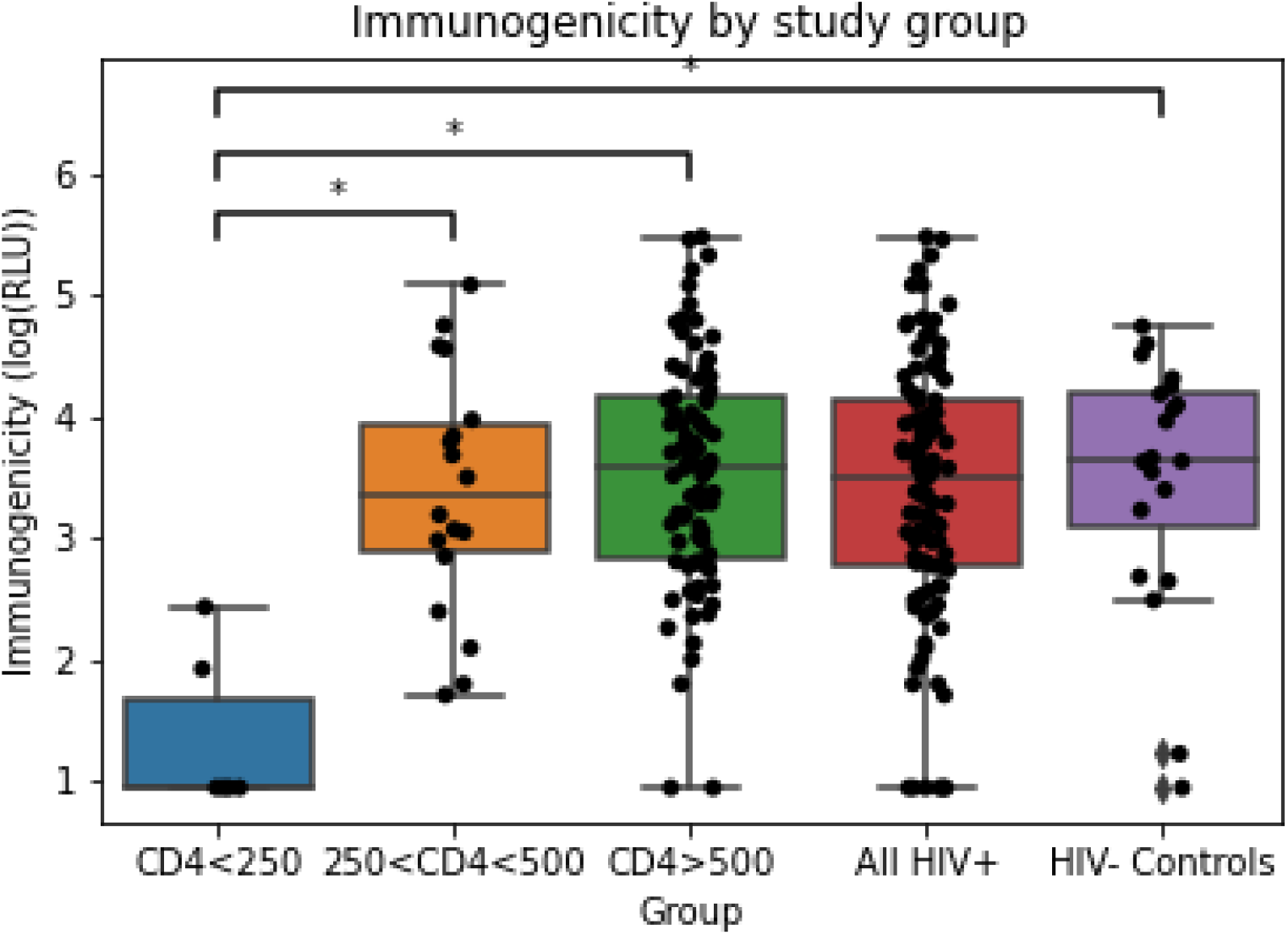
Immunogenicity in each study group. Immunogenicity (anti-RBD IgG response) was measured by ELISA and reported in RLU (relative luminescence units). RLU values log transformed for analysis. Statistically significant mean differences are denoted by * (Tukey test, p<0.001)

**Figure 2:**
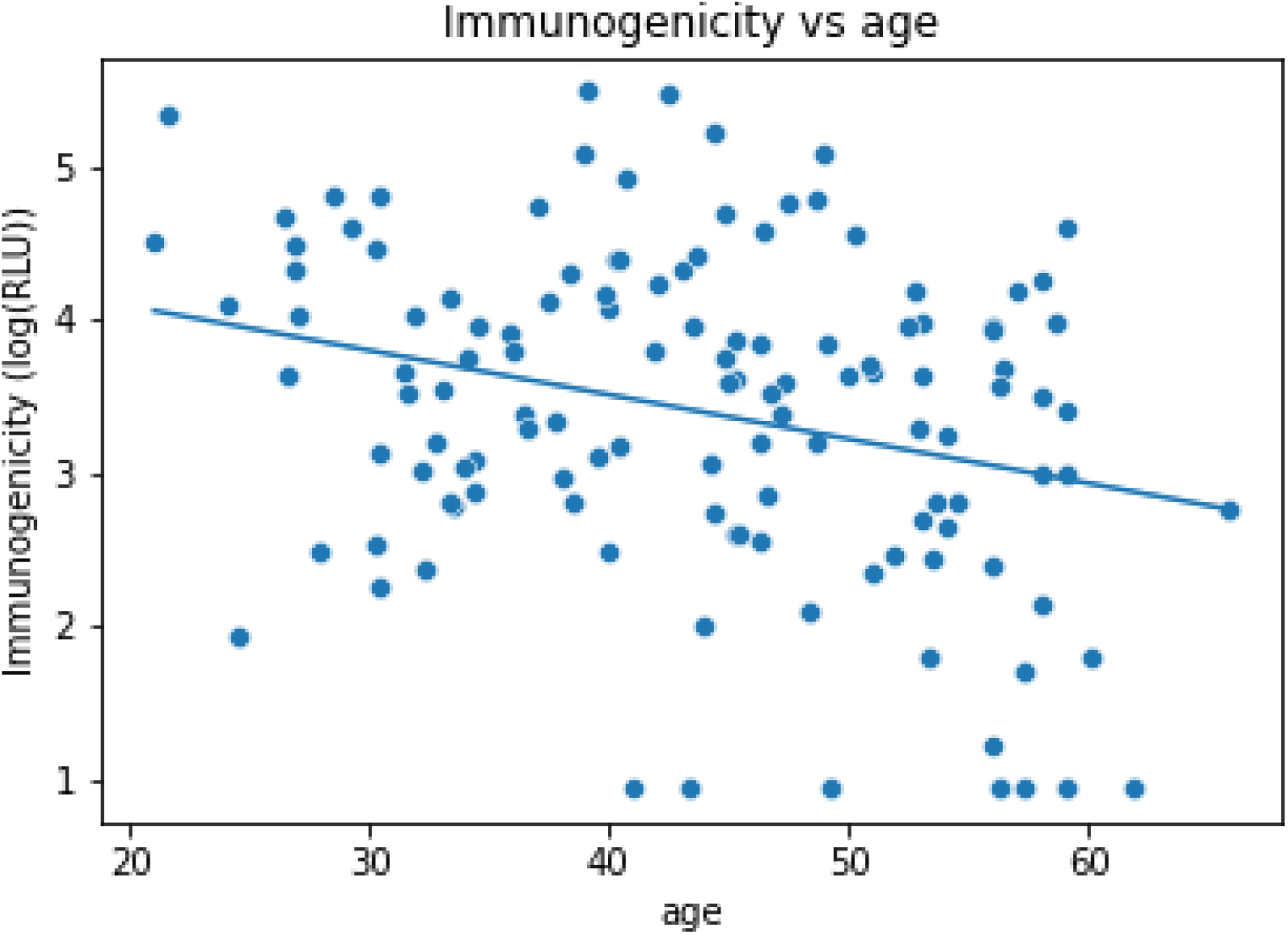
As part of the regression model, immunogenicity was found to be statistically significantly correlated with age (p<0.001). The magnitude of the association is weak, with an increase in 10 years corresponding to a decrease in 1.33 RLU (the range of ELISA RLU in this population was 2.56 (detection limit) to 236.03)

## Discussion

COVID-19 mRNA vaccines have been shown to be extremely efficacious in protecting against symptomatic disease, hospitalizations and death^1,2^. They were designed during the first wave of the pandemic when the main circulating strains were the original strain from Wuhan (D614) and the first variant of importance D614G^24^. Although these vaccines remain efficacious against most variants such as the Delta variant, a higher level of antibodies is required to confer optimal protection^25,26^. Immunocompromised populations are known to have weaker immune responses after vaccination. In the context of the emergence of variants which have a higher level of resistance to neutralization, it is important to ensure that this patient population mount an adequate response to vaccination. The level of vaccine immunogenicity may vary according to the type of immune deficiency. In recent studies evaluating immune responses to a COVID-19 vaccine in immunocompromised subjects, results varied according to the populations studied such as cancer patients^11^, transplant recipients^6,8^, hemodialysis patients^13,14^, or people treated with immunosuppressors^10^. This is expected as the immunosuppressive therapies used to treat these conditions target different pathways of the immune system, resulting in various degrees of impairment. In HIV-infected individuals, cellular immunity is mostly affected, CD4+ T lymphocytes being the target of this virus. CD4+ T cells are pivotal in orchestrating both the humoral and cellular immune responses to vaccination, and have an important impact on antibody production. In the past, it has been shown that people living with HIV-1 have lower responses to some types of vaccine and that these responses are dependent on the level of CD4+ T cells. With the development of more potent and well tolerated antiretrovirals to treat HIV-infection, a majority of people on treatment achieve an immune recovery with normalization of CD4 counts. However, even in this population, subtle defect in immune function persists^27,28^ and may impair vaccine response. Furthermore, a proportion of people living with HIV (PLWH) have very advanced disease with low CD4 counts and are at higher risk of not responding to vaccine. In our present study we show that if we look at the population of HIV-infected individuals as a whole, there is no significant difference in the level of anti-RBD IgG response as compared with a control group of HIV-negative individuals. However, when we stratify by CD4 counts, we see a statistically significant response between the groups, specifically between the group with CD4 below 250 cells/mm^3^ and the other groups. In pivotal clinical trials of COVID-19 vaccines, there was no statistically significant responses between different age groups. In our patient population, we do see an impact of age on immunogenicity after a single vaccine dose. Overall, our data show that some individuals in the lower CD4 cell stratum developed some responses to the vaccine, which support the hypothesis that this response could be increased by adding booster shots or modifying the dosing. While other published reports found no association between age and immune response to the vaccine in this population^15^, our data shows a statistically significant association after a single dose. These are preliminary data as these individuals will be followed over a one year period where we will be able to assess the durability and quality of these responses.

## Acknowledgments

This work was supported by le Ministère de l’Économie et de l’Innovation (MEI) du Québec, Programme de soutien aux organismes de recherche et d’innovation, the Fondation du CHUM, a CIHR foundation grant #352417, a CIHR operating grant Pandemic and Health Emergencies Research/Projet #465175 to A.F; an Exceptional Fund COVID-19 from the Canada Foundation for Innovation (CFI) #41027 to AF and DEK. A.F. is recipient of a Canada Research Chair on Retroviral Entry no. RCHS0235 950-232424. CT is the Pfizer/Université de Montréal chair on HIV translational research. V.M.L. is supported by a FRQS Junior 1 salary award. M.D. is supported by a FRQS Junior 2 salary award. D.E.K. is a FRQS Merit Research Scholar. LN is the recipient of a PREMIER fellowship. The funders had no role in study design, data collection and analysis, decision to publish, or preparation of the manuscript. We wish to thank Hélène Lanctor, Annie Chamberland, Stéphanie Matte, Étienne Larouche, Alla Belyansky, Ketsia Désormaux, Deborah Kleijnen, Tudor Luncean, Marc Messier-Peet, Chantal Morrisseau, Maude Vadeboncoeur, Sonia Yacini, Korotoum Wele Diallo, Morgane Guay-Jutras, Ouahiba Senouci, Mohamed Sylla, for their role in enrolling the subjects and processing the samples.

## References

1. Polack FP, Thomas SJ, Kitchin N, et al. Safety and Efficacy of the BNT162b2 mRNA Covid-19 Vaccine. N Engl J Med 2020;383:2603–15.

2. Baden LR, El Sahly HM, Essink B, et al. Efficacy and Safety of the mRNA-1273 SARS-CoV-2 Vaccine. N Engl J Med 2021;384:403–16.

3. Voysey M, Clemens SAC, Madhi SA, et al. Safety and efficacy of the ChAdOx1 nCoV-19 vaccine (AZD1222) against SARS-CoV-2: an interim analysis of four randomised controlled trials in Brazil, South Africa, and the UK. Lancet 2021;397:99–111.

4. Paris C, Perrin S, Hamonic S, et al. Effectiveness of mRNA-BNT162b2, mRNA-1273, and ChAdOx1 nCoV-19 vaccines against COVID-19 in health care workers: an observational study using surveillance data. Clin Microbiol Infect 2021.

5. Dagan N, Barda N, Kepten E, et al. BNT162b2 mRNA Covid-19 Vaccine in a Nationwide Mass Vaccination Setting. N Engl J Med 2021;384:1412–23.

6. Boyarsky BJ, Werbel WA, Avery RK, et al. Antibody Response to 2-Dose SARS-CoV-2 mRNA Vaccine Series in Solid Organ Transplant Recipients. JAMA 2021;325:2204–6.

7. Boyarsky BJ, Werbel WA, Avery RK, et al. Immunogenicity of a Single Dose of SARS-CoV-2 Messenger RNA Vaccine in Solid Organ Transplant Recipients. JAMA 2021;325:1784–6.

8. Marinaki S, Adamopoulos S, Degiannis D, et al. Immunogenicity of SARS-CoV-2 BNT162b2 vaccine in solid organ transplant recipients. Am J Transplant 2021.

9. Grupper A, Katchman H. Reduced humoral response to mRNA SARS-CoV-2 BNT162b2 vaccine in kidney transplant recipients without prior exposure to the virus: Not alarming, but should be taken gravely. Am J Transplant 2021.

10. Geisen UM, Berner DK, Tran F, et al. Immunogenicity and safety of anti-SARS-CoV-2 mRNA vaccines in patients with chronic inflammatory conditions and immunosuppressive therapy in a monocentric cohort. Ann Rheum Dis 2021.

11. Monin L, Laing AG, Munoz-Ruiz M, et al. Safety and immunogenicity of one versus two doses of the COVID-19 vaccine BNT162b2 for patients with cancer: interim analysis of a prospective observational study. Lancet Oncol 2021;22:765–78.

12. Bird S, Panopoulou A, Shea RL, et al. Response to first vaccination against SARS-CoV-2 in patients with multiple myeloma. Lancet Haematol 2021;8:e389–e92.

13. Anand S, Montez-Rath ME, Han J, et al. Antibody Response to COVID-19 vaccination in Patients Receiving Dialysis. medRxiv 2021.

14. Goupil R, Benlarbi M, Beaubien-Souligny W, et al. Short-term antibody response after 1 dose of BNT162b2 vaccine in patients receiving hemodialysis. CMAJ 2021;193:E793–E800.

15. Woldemeskel BA, Karaba AH, Garliss CC, et al. The BNT162b2 mRNA Vaccine Elicits Robust Humoral and Cellular Immune Responses in People Living with HIV. Clin Infect Dis 2021.

16. Touizer E, Alrubayyi A, Rees-Spear C, et al. Failure to seroconvert after two doses of BNT162b2 SARS-CoV-2 vaccine in a patient with uncontrolled HIV. Lancet HIV 2021;8:e317–e8.

17. Ruddy JA, Boyarsky BJ, Bailey JR, et al. Safety and antibody response to two-dose SARS-CoV-2 messenger RNA vaccination in persons with HIV. AIDS 2021.

18. Frater J, Ewer KJ, Ogbe A, et al. Safety and immunogenicity of the ChAdOx1 nCoV-19 (AZD1222) vaccine against SARS-CoV-2 in HIV infection: a single-arm substudy of a phase 2/3 clinical trial. Lancet HIV 2021.

19. Beaudoin-Bussieres G, Laumaea A, Anand SP, et al. Decline of Humoral Responses against SARS-CoV-2 Spike in Convalescent Individuals. mBio 2020;11.

20. Anand SP, Prevost J, Nayrac M, et al. Longitudinal analysis of humoral immunity against SARS-CoV-2 Spike in convalescent individuals up to 8 months post-symptom onset. bioRxiv 2021.

21. Gasser R, Cloutier M, Prevost J, et al. Major role of IgM in the neutralizing activity of convalescent plasma against SARS-CoV-2. Cell Rep 2021;34:108790.

22. Prevost J, Gasser R, Beaudoin-Bussieres G, et al. Cross-Sectional Evaluation of Humoral Responses against SARS-CoV-2 Spike. Cell Rep Med 2020;1:100126.

23. Tauzin A, Nayrac M, Benlarbi M, et al. A single dose of the SARS-CoV-2 vaccine BNT162b2 elicits Fc-mediated antibody effector functions and T cell responses. Cell Host Microbe 2021;29:1137–50 e6.

24. Zhang J, Cai Y, Xiao T, et al. Structural impact on SARS-CoV-2 spike protein by D614G substitution. Science 2021;372:525–30.

25. Lopez Bernal J, Andrews N, Gower C, et al. Effectiveness of Covid-19 Vaccines against the B.1.617.2 (Delta) Variant. N Engl J Med 2021.

26. Planas D, Veyer D, Baidaliuk A, et al. Reduced sensitivity of SARS-CoV-2 variant Delta to antibody neutralization. Nature 2021.

27. Morou A, Brunet-Ratnasingham E, Dube M, et al. Altered differentiation is central to HIV-specific CD4(+) T cell dysfunction in progressive disease. Nat Immunol 2019;20:1059–70.

28. Niessl J, Baxter AE, Morou A, et al. Persistent expansion and Th1-like skewing of HIV-specific circulating T follicular helper cells during antiretroviral therapy. EBioMedicine 2020;54:102727.

